# Discovery of Latent Drivers from Double Mutations in Pan-Cancer Data Reveal their Clinical Impact

**DOI:** 10.1101/2021.04.02.438239

**Authors:** Bengi Ruken Yavuz, Chung-Jung Tsai, Ruth Nussinov, Nurcan Tuncbag

## Abstract

**Background**

Transforming patient-specific molecular data into clinical decisions is fundamental to personalized medicine. Despite massive advancements in cancer genomics, to date driver mutations whose frequencies are low, and their observable transformation potential is minor have escaped identification. Yet, when paired with other mutations *in cis*, such ‘latent driver’ mutations can drive cancer. Here, we discover potential ‘latent driver’ double mutations.

**Method**

We applied a statistical approach to identify significantly co-occurring mutations in the pan-cancer data of mutation profiles of ∼80,000 tumor sequences from the TCGA and AACR GENIE databases. The components of same gene doublets were assessed as potential latent drivers. We merged the analysis of the significant double mutations with drug response data of cell lines and patient derived xenografts (PDXs). This allowed us to link the potential impact of double mutations to clinical information and discover signatures for some cancer types.

**Results**

Our comprehensive statistical analysis identified 228 same gene double mutations of which 113 mutations are cataloged as latent drivers. Oncogenic activation of a protein can be through either single or multiple independent mechanisms of action. Combinations of a driver mutation with either a driver, a weak driver, or a strong latent driver have the potential of a single gene leading to a fully activated state and high drug response rate. Tumor suppressors require higher mutational load to coincide with double mutations compared to oncogenes which implies their relative robustness to losing their functions. Evaluation of the response of cell lines and patient-derived xenograft data to drug treatment indicate that in certain genes double mutations can increase oncogenic activity, hence a better drug response (e.g. in PIK3CA), or they can promote resistance to the drugs (e.g. in EGFR).

**Conclusion**

Our comprehensive analysis of same allele double mutations in cancer genome landscapes emphasizes that interrogation of big genomic data and integration with the results of large-scale small-molecule sensitivity data can provide deep patterns that are rare; but can still result in dramatic phenotypic alterations, and provide clinical signatures for some cancer types.

## Background

Cancer is a disease of uncontrolled cell proliferation driven by molecular alterations. The impact of these alterations diffuses into the molecular interaction network and changes signaling pathways and transcriptional regulation in the cell. Not all alterations equally contribute to growth advantage of cancer cells. Some mutations are drivers; others are passengers [1]. Whereas it is generally believed that passenger mutations do not bestow proliferative effects on the disease phenotype, their properties and possible roles are not fully understood [2]. Comprehensive screening of thousands of p53 mutations and phenotypic characterization of these mutations have shown that mutations that maintain wild-type functionality of p53 are unlikely to be cancer drivers [3]. However, cancer genomics and evolutionary studies suggest that the accumulation of ‘slightly’ deleterious passenger mutations can slow cancer progression and this could be exploited for therapeutic purposes [4]. Lately, another class of mutations was defined, dubbed “latent” or “mini-drivers” [5–7]. Latent mutations may assume a driver-like behavior yet were not identified as drivers per se. Latent drivers emerge during cancer evolution and their detection may help forecast cancer progression and improve personalized treatment strategies [6]. Driver mutations are classified into three types, strong driver, driver and weak driver. As for latent drivers, there are strong latent and weak latent drivers. Curated driver genes and mutations have been deposited in multiple databases [8–10]and used by multiple research groups to develop computational approaches to predict driver genes and driver mutations [11–16]. These methods, including the frequency-based methods, subnetwork identification methods, and 3D mutation search methods, have been comprehensively compared [17–19]. One of the concerns with frequency-based approaches is that prohibitively large sample sizes are needed to identify infrequently mutated driver genes. Thus, in frequency-based approaches, there is a risk of generating biased results due to background mutation rates [20]. Large databases catalog cancer driver genes and driver mutations and help in understanding the mechanism behind tumorigenesis. However, frequency-based approaches fail in the identification of rare drivers which can be tissue-specific [21]. A recent multidimensional analysis of cancer driver genes in IntOGen showed that some drivers are cancer-wide whereas others are specific to a limited number of cancer types [14].

Even a single mutation in a gene can be considered as a prognostic marker and change the global genome and protein expression, eventually altering the signaling pathways [22]. However, it has been estimated that the contribution of a single driver mutation to cancer progression is very small and needs additional mutations over time [23]. Despite DNA repair, somatic mutations accumulate and different genotypes in individual tissues are generated. This mechanism, called ‘somatic mosaicism’, offers driver or synergistic mutations an advantage in cancer cells [24].

Recently, the combination of single frequent mutations with a rare, or weak mutation in the *same* gene was shown to have a significant advantage in tumor progression and influence treatment response. These double mutations *in cis* in PIK3CA were shown to be more oncogenic, and more sensitive to an inhibitor compared to a single mutation [25]. A recent work cataloged ‘composite mutations’ of *multiple* genes – i.e. acting through same proteins – having more than one non-synonymous mutation in the same tumor [26]. Saito et al demonstrated the functional implications of multiple driver mutations in the same oncogene with an emphasis on PIK3CA [27]. Analysis of the rare mutations in cancer patients revealed known and hidden onco-drivers that are mutually exclusive in the same pathway suggesting epistatic mechanisms [28]. Many approaches are based on the principle that functionally-related genes have similar profiles of epistatic interactions [29]. One proposed explanation, typically for mutations in the same cellular pathway, involves functional redundancy. After a pathway has been mutated once, there is no evolutionary benefit to the clone from additional mutations in that pathway [29, 30].

Here, aided by informatics techniques, we systematically screen somatic mutations in pan-cancer data across ∼80,000 patient tumors. We aim to find co-occurring patterns that are predominantly present in specific tissues and tumor types. Our screening reveals tumor-type specific double mutations on the same gene which may promote tumorigenesis and alter the response to treatments. It also reveals that tumors having at least one double mutations pair can lead to changes in response to drugs. We cataloged the components of double-mutations as latent mutations if their co-occurrence is significant and not yet labeled as a cancer driver. This led us to uncover 113 latent driver mutations. The oncogenic activation of a gene is through either single or multiple independent mechanisms of action. We present these different mechanisms through the same gene double mutations. Although the existence of a set of driver genes is considered cancer-wide, we show that having double mutations on those genes is cancer-specific. Same gene double mutations are relatively rare; however, their impact is elevated in tumor progression.

## Methods

### Data collection and Processing

All available somatic missense mutation profiles are downloaded from two sources, The Cancer Genome Atlas (TCGA) and the AACR launched Project GENIE (Genomics Evidence Neoplasia Information Exchange) [31–33]. The TCGA mutation annotation file contains more than 11,000 human tumors across 33 different cancer types. The GENIE mutation file (Release 6.2-public) contains 70679 samples across 671 cancer subtypes under Oncotree classification. The GENIE cohort contains multiple tumor barcodes belonging to the same tumor type. In such a case only one primary tumor barcode is kept for further analysis. We continued the analysis with 78837 samples from 671 cancer subtypes and 34 tissues (including UNKNOWN and OTHER categories).

### Identification of Significant Double Alterations

The total number of mutations is 1638191 in 19443 genes. We only evaluated dual combinations of 21983 (on 5062 genes) of these alterations observed on at least 5 tumors and constructed binary combinations of them. Then we created a contingency table for each combination of tumor numbers having both alterations, only the first or second alteration and none of those two alterations. Based on the contingency table, we calculated the p-value by using Fisher Exact Test with the formula below:

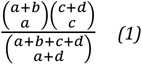

where a is the number of tumors having both alterations, b is the number of tumors having only the first alteration, c is the number of tumors having only the second alteration and d is the number of tumors not having these two alterations.

228 significant pairs and 227 non-significant pairs were decided using the Fisher Exact Test for p=0.05. We used the Catalog of Validated Oncogenic Mutations from the Cancer Genome Interpreter [10] to label dual mutation components: if a mutation is among the 5601 driver mutations, we label it as known driver (D), otherwise potential latent driver (d). We then classified a known driver mutation as a driver if it is present in more than 500 tumors; otherwise, it is a weak driver. Similarly, we dubbed a potential latent driver mutation as a strong latent driver if it is present in more than 10 tumors; otherwise, we classified it as a weak latent driver.

Additionally, double mutations are annotated based on their functions, domains, chemical properties and structural proximity (see Supplementary Text)

### Survival Analysis

For survival analysis, 10336 patients in MSK impact 2017 and 11160 patients in TCGA and their overall survival status are used [31, 33]. We compared survival times of tumor groups with significant same/different gene double mutations and single mutations in a specific cancer subtype. The first group is the union of patients with significant doublets whereas the second is the union of patients that carry only one component of these significant double mutations. Then we gathered overall survival times (time in months) and vital status (1: Deceased, 0: Alive) of these patients for survival analysis.

We utilized the “survival” library of R to do Kaplan Meier Survival Analysis of double and single mutant groups. The survival probability at any particular time is calculated by the formula given below [34]:

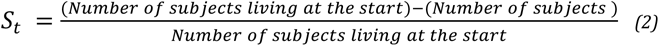

### Oncoprint Maps

To reveal mutual exclusivity and co-occurrence patterns between double mutations we plotted oncoprint maps by using ComplexHeatmap package of R [35].

### Cell Line Network Construction

We obtained a list of cell lines with the dual mutations from Cell Model Passports and their drug response information from CancerrxGene [36, 37]. We also extracted information about drug targets and target pathways. We used 2 different approaches to select drugs for PTEN, APC, and PIK3CA dual mutant cell lines: if a drug is in the gray zone (|z-score|<=2) in the single mutant cell lines but gives a significant drug response in a dual mutant cell line (|z-score|>2). If there is a single mutant cell line that is sensitive (or resistant) to the drug but the dual mutant cell line gives an opposite response to the drug. (Drug response flips sensitive into resistant or resistant into sensitive between single and dual mutant cell lines).

For EGFR we selected drugs that give significant drug response either in the single or dual mutant cell line. Then we formed networks connecting mutations to cell lines, cell lines to drugs, and drugs to their target pathways.

### Patient-Derived Xenograft Analysis

We used the mutation profiles, transcriptomic data and drug responses of patient-derived xenografts in [38]. We determined xenografts harboring significant doublets. Then, we compared changes in tumor volumes of single and dual mutant xenografts for the untreated and drug-treated cases (single mutation is part of a significant dual mutation). We preferred to specify the time intervals in multiples of 5. When a given timepoint is not a multiple of 5, we used linear interpolation between two nearest numbers containing a multiple of 5.

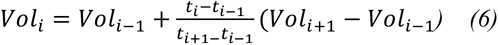

where t_i_ is a timepoint that is multiple of 5 between the given timepoints t_i-1_ and t_i+1_ and Vol_i_ is the volume (mm^3^) at timepoint i.

## Results

### Discovery of Latent Drivers through Double Mutations

The availability of a vast amount of pan-cancer genomic data helps to find mutational patterns that can be signatures of the specific tumor tissues or cancer types. Multiple mutations in a single gene rarely co-occur in patient tumors. However, when they are together, they may cause dramatic phenotypic differences [25–27]. For example, dual mutations in PIK3CA increase the sensitivity to PI3K inhibitors in breast cancer [25], while dual mutations in EGFR predominantly exist in lung cancer [39]. A strong driver may couple with a weak driver or a latent driver to increase the pathological impact of the alterations. This pattern also gives insight into the latent drivers that are context specific. We exploited the dual mutations to discover latent drivers. For this purpose, following the Oncotree classification we obtained and cataloged missense mutation profiles of ∼80,000 tumors from TCGA and GENIE Pan-Cancer datasets from 34 main tissues and 672 cancer subtypes including tissues tagged as Unknown and Other (Figure 1A). Collecting all missense mutations on each gene and counting their pairwise combinations result in 228 significant double mutations (p-value < 0.05, Fisher exact test). Especially, when single mutations across patient tumors are systematically reduced in the co-occurring mutation patterns, the double mutations are revealed to be cancer specific. We also assembled tissue-specific sets of double alterations since tissues differ in sample size and are enriched in different genes and mutations. As shown in Figure 2A, co-occurring double mutations on the same gene are relatively rare, with varied frequencies across tissues. In some cancer tissues, doublets are present on the same gene in up to 10% of the patient tumors. However, same gene doublets are either extremely rare or not present in other tissues, such as the pancreas, ovary, liver, kidney, biliary tract. Same gene double mutations accumulate on 35 genes in the pan-cancer dataset of which 20 genes are tumor suppressors (TSG), 12 are oncogenes (OG) and the rest labeled as both.

**Figure 1.**
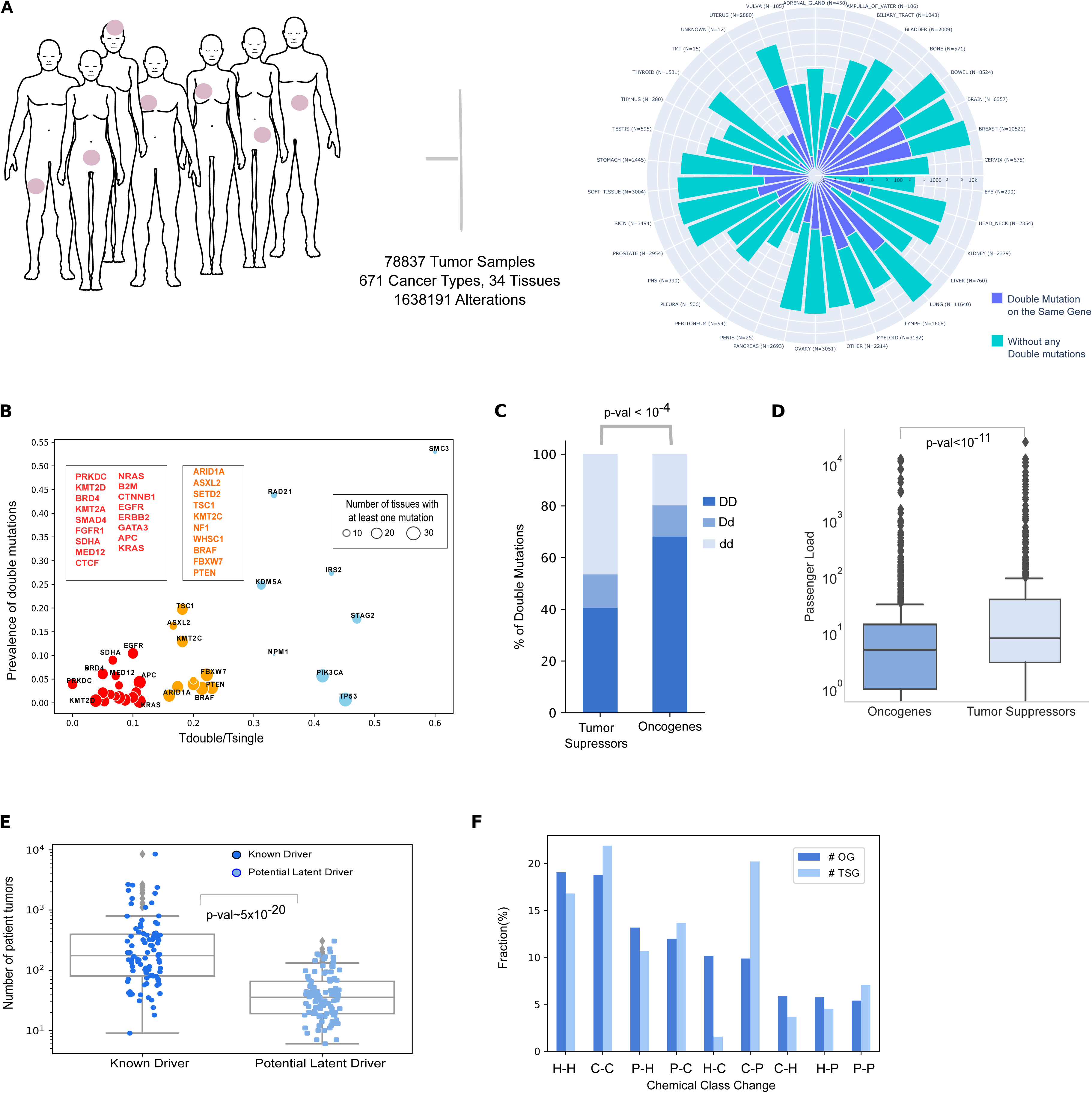
Overall statistics of the data set, mutation load and double mutations, and analysis of the significant double mutations. **(A)** Total number of tumors, alterations, cancer types in the union of TCGA and AACR GENIE studies. Same gene double and different gene double mutations are found by the Mann-Whitney U test. Windrose plot showing the number of same gene double mutant (blue) tumors across 34 tissues (Oncotree) on the log-scale axis. Green portion represents the amount of tumors without any significant double mutation. **(B)** Tissue specificity of same gene dual mutations compared to their single mutant counterparts. X-axis shows the ratio of the number of tissues containing double and single mutant tumors in each gene. Y-axis shows the fraction of overall count of double mutant tumors to the single mutant ones. Smaller values along the x-axis indicates tissue specific same gene double mutations. Genes having cancer-specific double mutations are red and cancer-wide double mutations are in blue. **(C)** Composition of the double mutations based on known driver (D) and potential latent driver (d) labels in tumor suppressor genes and oncogenes where D is already known frequent driver mutations, d is relatively rare potential latent drivers. Fraction (%) of DD, Dd, dd type double mutations are significantly different between oncogenes and tumor suppressor genes. **(D)** Box plot showing passenger mutation load in tumor suppressor genes and oncogenes. The patient group carrying same gene double mutations on oncogenes have relatively smaller passenger mutation counts compared to the group carrying double mutation on tumor suppressor genes. **(E)** Tumor count distributions of known driver and potential latent driver mutations. Known driver mutations are observed more frequently that the potential latent driver mutations. **(F)** Grouped bar plot shows the fraction (%) of alterations in chemical properties of amino acids for oncogenes and tumor suppressor genes. Mutations on oncogenes mostly convert hydrophobic residues to charged, and mutations on tumor suppressor genes usually convert charged residues to polar.

**Figure 2.**
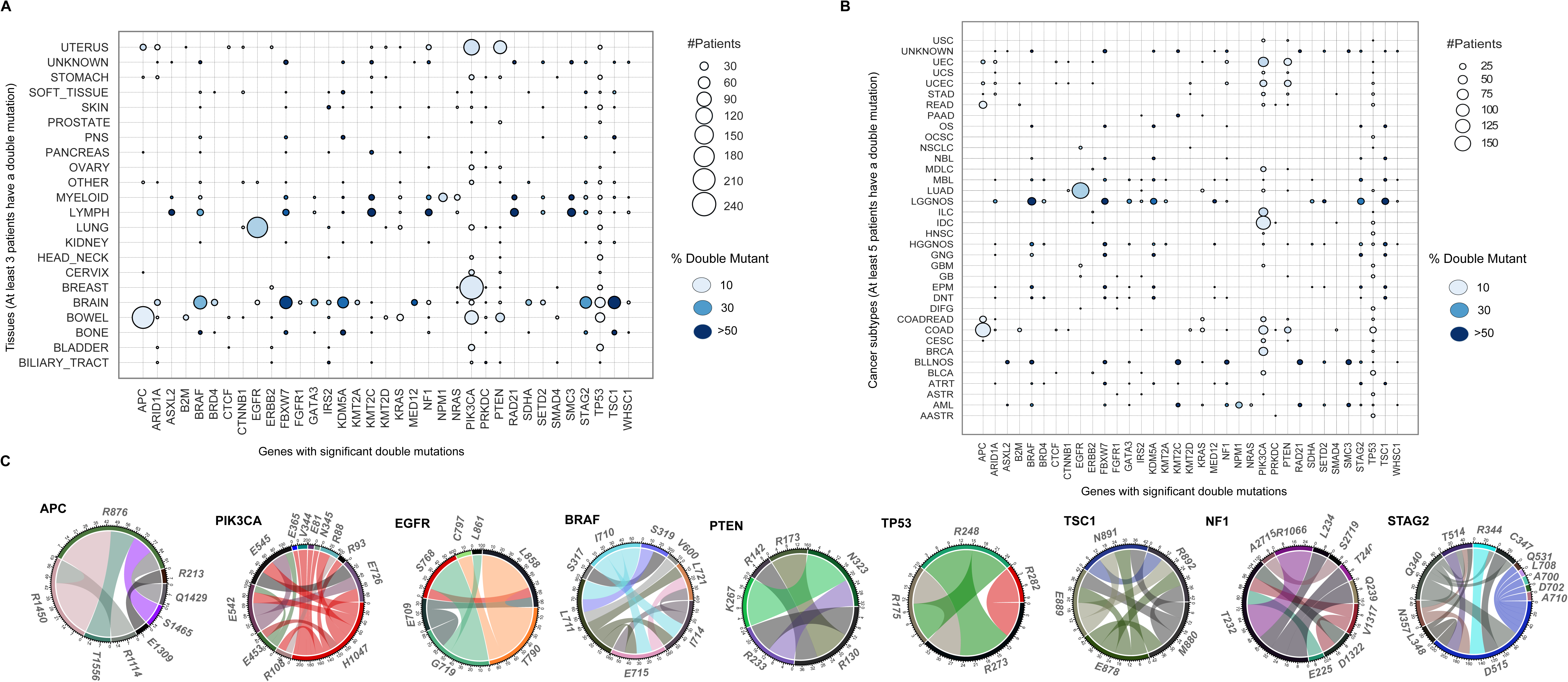
Same gene double mutations are specific to some tissues or cancer subtypes. Bubble plots show number (node size) and frequency (node color) of double-mutant tumors among gene-mutant tumors across different tissues and cancer subtypes (Oncotree). For the 35 genes with significant same gene double mutations, node size represents the number of patients carrying at least one doublet on a gene in a tissue or cancer type. **(A)** Presence of same gene double mutations across different cancer tissues where at least 3 tumors carry at least one same gene double mutation on one of the 35 genes. **(B)** Presence of different gene dual mutations across different cancer subtypes. The cancer subtypes where at least 5 tumors carry at least one double mutation are listed on the y-axis. **(C)** Representation of mutations in genes to compose a doublet as a circular diagram. Circumference of the circles divided into arcs proportional to the frequency of each mutated residue. The strips from one residue to another represents significant double mutations with size of strips indicating frequency of each double mutation.

Recently, the frequency of driver genes was analyzed together with the maximum prevalence of their mutations, distinguishing cancer-specific drivers versus cancer-wide drivers [14]. We applied a similar analysis to our dataset composed of double mutations on the same gene where we obtained the ratio of the number of tissues carrying double mutations (T_double_) and single mutations (T_single_). We also calculated the prevalence of double mutations compared to single mutations. As a result, although some genes and their single mutant states have been previously cataloged cancer-wide, we found sets of double mutations that are cancer tissue-specific. Examples include double mutations in PTEN, EGFR, and KRAS (Figure 1B).

We retrieved the known driver mutations from the Cancer Genome Interpreter database to evaluate if the double mutations are composed of known drivers or other mutations that are not cataloged as drivers but can be considered as ‘potential latent driver’ mutations. In a doublet, the components can be known drivers or potential latent drivers, so each doublet is cataloged as DD, Dd and dd. That is, DD is a known driver-known driver doublet, Dd is a known driver-potential latent driver and dd is a doublet consisting of two potential latent drivers. Among the 228 same gene double mutations, there are 115 DD, 28 Dd, 85 dd where the mutations that are not catalogued as driver are potential latent drivers (the 228 same gene double mutations are composed of 91 known major drivers, 113 potential latent drivers). Thus, our analysis can capture rare mutations that are potential latent driver candidates. We observe that oncogenes have significantly more DD mutations than tumor suppressors (p-value < 10^-4^), although their background probability to have a double mutation is similar (Figure 1C). This result implies that becoming more oncogenic requires mostly co-occurrence of two frequent mutations while suspending tumor suppressor activities may involve rare mutations coming together.

Tumor suppressor genes have 131 double mutations in 883 patient tumors and oncogenes have 91 double mutations in 1000 patient tumors. Patient tumors that have at least one double mutation in any TSG have a significantly higher passenger mutation load compared to patient tumors having at least one double mutation in an oncogene (p-value < 10^-11^, Figure 1D). These results imply that double mutations are very rare. Especially tumor suppressor genes require a very high mutation load for two coexisting mutations in a single gene. Based on the mutation load, and in line with our previous result, loss of function through double mutations in TSGs requires considerably higher mutational load compared to gain of function in oncogenes.

Known driver mutations have a higher frequency than potential latent driver mutations (Figure 1E). The median values of tumor counts for known driver and potential latent driver mutations are 170 and 35.5, respectively (p-value < 5×10^-20^). Potential driver mutations are relatively rare, and their pathological impact can be dramatic when they couple with another mutation. Therefore, we cataloged all potential latent driver mutations that contribute to a significant doublet in the same gene as strong or weak latent drivers. The list of 113 latent drivers is given in Table S1.

Next, we followed a bottom-up approach to obtain the spatial, chemical, and pathway level organization of the double mutations. We used the pan-cancer mutation clusters deposited in 3DHotspot where each cluster represents the set of mutations that are spatially close to each other [40]. We found that components of the doublets in the same gene are usually spatially distant from each other. The simultaneous presence of spatially close two strong driver mutations is very rare in a patient tumor. However, some weak drivers are proximal to either a strong driver or another weak driver, as in the cases of mutations at positions 711, 714, 715 in BRAF. Spatially close residues may form potent allosteric couples, which may enhance proliferation.

Analysis of the chemical class of doublets in oncogenes and tumor suppressor genes harboring the same gene doublets revealed that Charged-Polar and Hydrophobic-Charged switches are more dominant among tumor suppressors and oncogenes respectively (Figure 1F). Double mutations are either located in flexible or hinge or disordered regions or in different domains (Supplementary Text, Figure S1A). Components of each double are annotated in different molecular functions (Supplementary Text, Figure S1B).

### Doublets on the Same Gene are Rare, but are a Signature for Some Cancer Types

Figure 2A illustrates the tissue-specific prevalence of double mutations in the same gene. TP53 and its double mutations are cancer wide. A recent study verified that PIK3CA double mutations *in cis* increase oncogenicity and sensitivity to PI3Kα inhibitors [25]. In our dataset, PIK3CA double mutations are also quite common in breast and uterus tumors. In the uterus, PIK3CA mutations are more inclined to constitute double mutations (around 90%). Among lung tumors, EGFR and bowel tumors APC double mutations are ahead by far. Bowel, breast, and lung tissues are enriched with double mutations on specific genes whereas brain tissue has significant double mutations in multiple genes such as BRAF, FBXW7, KDM5A, STAG2, TP53, TSC1. LUAD (Lung Adenocarcinoma) is enriched with EGFR dual mutations. 90% of the EGFR mutations are in more than 160 tumors. COAD (Colon Adenocarcinoma) is enriched with APC and PTEN dual mutations. We note that PIK3CA double mutations are relatively more dominant in BRCA, COAD, and UCEC subtypes (Figure 2B). A set of known driver mutations, for example in KRAS and IDH1 are usually present as single mutations but are frequently paired with mutations in other genes. The most frequent IDH1 mutation occurs at position 132 located in the interface of its homodimer [41].

The most frequent mutation, KRAS^G12D^, is rarely coupled with another mutation in KRAS. The mutational mosaic of KRAS is distinguishable in different cancer types. KRAS^G12D^ is predominantly present in pancreatic and colorectal cancers [42]. KRAS mutations are context-specific and a mutation may act in different cancers. However, among this limited number of KRAS double mutations, KRAS^G12D/A149T^ accumulates in lung tissue and only exists in primary tumors in our dataset. KRAS^A146T^ promotes opening of Switch I in GEF mediated GDP-GTP nucleotide exchange whereas KRAS^G12D^ abolishes GAP-mediated hydrolysis [43].

Figure 2C illustrates some sequence details. In APC, EGFR, PTEN, and TP53 the diversity of the double mutations is limited, but this is not the case in PIK3CA and BRAF. Among them, BRAF^V600E^, a strong driver, is rarely coupled with another BRAF mutation, but the rest are (Figure S2A). Other BRAF mutations such as at 711, 714, 715, and 721 are close to each other and coupled in a set of patients, especially in brain tissue (Figure S2B and S2C).

Another interesting case is the double mutations in the cohesin complex. Mutated cohesin can enhance Wnt signaling by stabilizing beta-catenin [44]. Targeting Wnt signaling in cohesin mutant cancer cells was proposed as a novel therapeutic strategy. Double, even multiple mutations in the components of the cohesin complex (Figure S3A) in the same tumor may dramatically increase Wnt signaling. In Figure S3B, we notice that STAG2 double mutations accumulate in the brain, RAD21 and SMC3 double mutations are prominent in lymph and myeloid tissues.

### The Clinical Impact of Double Mutations in PIK3CA

PIK3CA is a large protein with drivers e.g. H1047R, E545K and weak drivers such as R88Q, E453K, M1043I. It is the second (or third) most highly mutated protein and its number of double mutations is also relatively higher than other proteins. The pathological impact of a single driver may be insufficient. Full activation of oncogenic PIK3CA is through two drivers acting in different, albeit complementary mechanisms. One well-known example is H1047 and E545 double mutations enhancing proliferation [45]. However, E545 and E542 double mutations do not make PIK3CA reach the fully activated level. Also, the combination of two strong latent driver mutations – but not two weak – can act like a driver mutation.

In our dataset of significant double mutations, 46 of the 228 are in PIK3CA. Enhanced activation of PIK3CA via dual mutations is shown in Figure 3A, where most of them are composed of one frequent and one rare mutation. Our frequency-based analysis revealed that P104, E726 and M1004 might be a strong latent drivers coupled with a driver mutation. PIK3CA double mutations are also tissue- and context-specific as shown in Figure 3B. Most are in breast tissue. An exception involves R88Q doublets which are depleted in breast but frequent in uterus tumors. Their structural location is shown in Figure 3C. Kinase mutations work by destabilizing the inactive or stabilizing the active state. These are better captured by their detailed conformational consequences. The mechanisms of activation of PI3Kα by these driver mutations have been recently worked out [45–47]. Unsurprisingly, considering their diverse mechanisms of action no clear trend is observed in the calculated folding free energy (ΔΔG) upon double or single mutation with DynaMut [48] (Suppl. Text and Figure S4). If the components of double mutations act via distinct mechanisms, the additivity of their activation potential is high; otherwise the additivity is low as in the E545/E542 example where the mutations execute the same mechanism of action.

**Figure 3.**
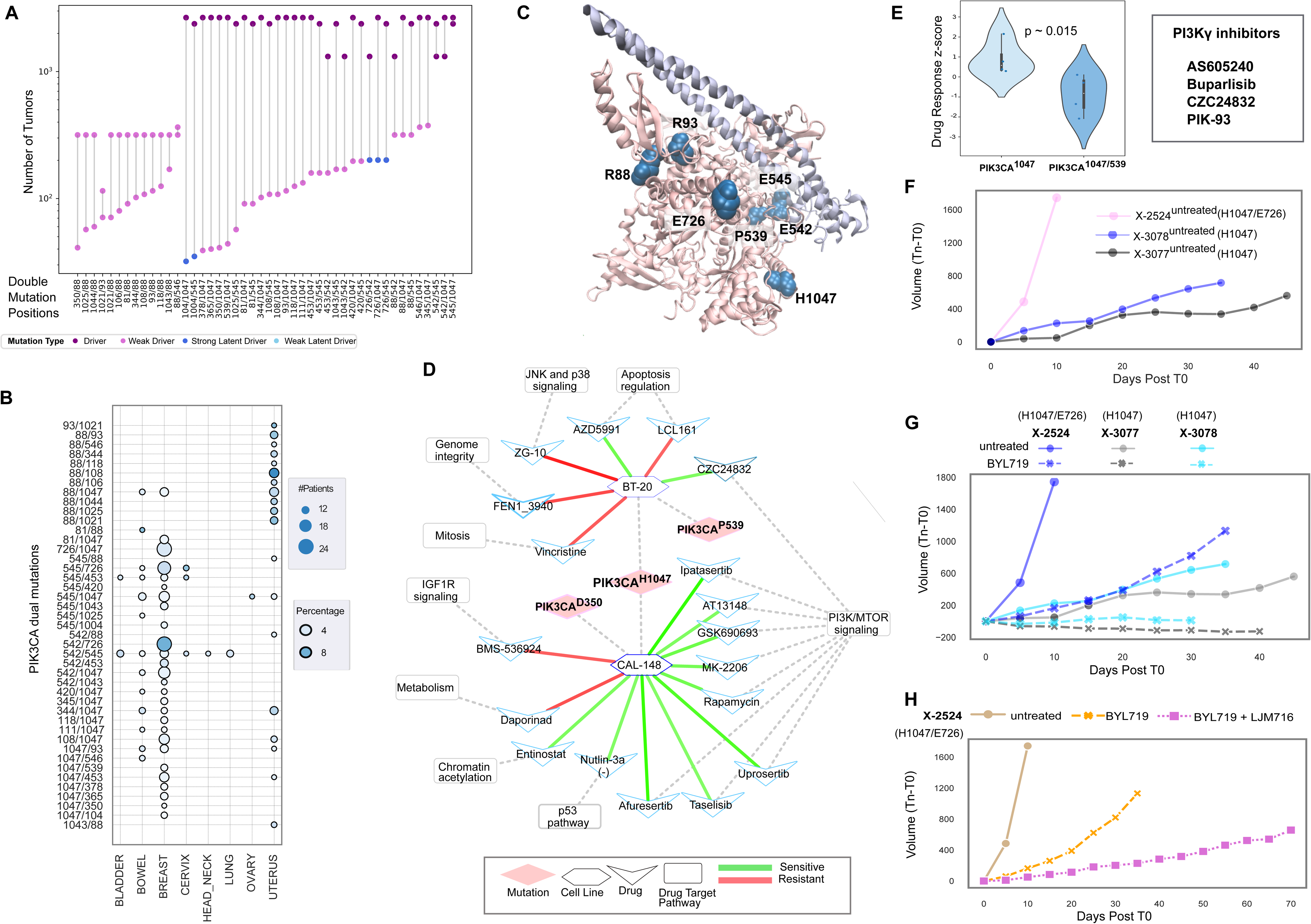
A detailed analysis of PIK3CA double mutation profile, 3D structure and clinical implications. **(A)** Paired dot plot of the 46 double mutations on PIK3CA, and the number of tumors carrying them. Colors indicate type of a mutation, driver (purple), weak driver (orchid), strong latent driver (blue), weak latent driver (light blue). Drivers and weak drivers are known driver mutations with ≥500 and <500 harboring tumors respectively. Strong latent drivers and weak latent drivers are potential latent driver mutations carried by ≥10 and <10 tumors respectively. There are 3 *driver/driver*, 13 weak *driver/weak driver*, 5 *driver/strong latent driver,* 35 *driver/weak driver. P104, E726 and M1004 have a potential to be strong latent drivers.* **(B)** Presence of PIK3CA same gene dual mutations across different cancer tissues. Dots are scaled based on the number of tumor having double mutations, and color corresponds to the percentage of double mutant tumors among single mutants. **(C)** 3D structure of PIK3CA (PDB: 4OVV) with H1047, E726, E542, E545, R88, R93, P539. **(D)** Response of PIK3CA double mutant breast cancer cell lines to drugs in network representation. BT-20 (H1047/P539) and CAL148 (H1047/D350) becomes sensitive to PI3K/MTOR pathway targeting drugs. **(E)** PIK3CA mutation doublets in breast cancer and the associated violin plot illustrating response to PI3Kɣ inhibitors. H1047/P539 double mutant tumor becomes more sensitive to PI3Kɣ inhibitors **(F)** Tumor volume change of single and double PIK3CA mutant xenografts without any treatment. There is a remarkable tumor increase in the double mutant xenograft. **(G)** Tumor volume comparison of the single and double mutant xenografts without any treatment and with BYL719 (Alpelisib) treatment, the double mutant xenograft responds better to the PI3K⍺ inhibitor drug. **(H)** Comparing tumor volume changes of the dual PIK3CA mutant xenografts without any treatment and with BYL719 and BYL719+LJM716 treatment. The tumor growth of the double mutant xenograft X-2524 is prominently slow when a combination therapy BYL719+LJM716 is applied compared to untreated and BYL719 treatment alone.

The impact of co-occurring mutations in the same gene is mostly additive but can be also cooperative. There are seven allosteric mutations at positions 83, 88, 365, 539, 542, 603, 629 in PIK3CA in BRCA as cataloged in Allosteric DB [49]. We found 23 significant double mutations in PIK3CA in the BRCA subtype of breast tumors. Dual mutations PIK3CA^1047/88^, PIK3CA^1047/539^, PIK3CA^1047/539^ are composed of one known driver (at position 1047) and one weak driver mutation (PIK3CA^88^ and PIK3CA^539^) which are allosteric mutations. Their effects are additive.

To further evaluate the double mutations, we used cancer cell lines from the DepMap project and patient-derived xenograft (PDX) samples in [38]. In both datasets, mutation profiles and response to drug treatment information are available. Additionally, we used the temporal data on tumor volume growth in PDX samples in untreated conditions and drug-treated conditions.

We found two breast cancer cell lines belonging to the BRCA subtype: BT-20 has a double mutation PIK3CA^1047/539^ and Cal-148 has PIK3CA^1047/350^. H1047R is a frequent driver. However, 539 and 350 are rare mutations in the Pan-cancer data, making them weak drivers. To explore the impact of the double mutations in terms of drug response, a network of cell lines to drugs and target pathways is constructed (Figure 3D) where drugs are linked to each cell line which has altered response compared to their single mutation counterparts. Cal-148, which has PIK3CA^1047/350^, is more sensitive to drugs targeting the PI3K/mTOR pathway compared to the single mutant cell lines. Indeed, we found a difference in the response to PIK3α inhibitors in double-mutant cell line BT-20 which is more sensitive to this class of inhibitors compared to single mutant cell line counterparts (p-value=0.015). Additionally, an evaluation of other classes of inhibitors showed that the PIK3K inhibitor CZC24832 does not work on single mutant MFM-223, but double mutant BT-20 is sensitive to it (Figure 3E).

We retrieved PDXs having double mutations in PIK3CA to explore the tumor volume changes and drug responses compared to single mutant PDXs. Properties of the patient tumors can be maintained in xenografts and can help assess the impact of double mutations. We found three PDXs having double PIK3CA mutations (726/1047, 88/542, 88/1025). In PDX X-2524 H1047R/E726K, a strong driver/strong latent driver combination, the volume change of the tumor between days 0 and 10 is more than 1700 mm^3^, while single mutant tumors X-3077 and X-3078 (with mutation H1047R) have volume change of ∼200 mm^3^ in the first 10 days reaching ∼400 mm^3^ at around 35 days (Figure 3F). H1047R/E726K tumors grow significantly faster compared to the single mutant case.

We analyzed the effects of drugs on tumor growth of these three PDX tumors. We observed that BYL-719 (Alpelisib), a selective PI3Kα inhibitor, diminishes tumor volume by 88% (around 1600 mm^3^) in the first 10 days in the double mutant in the xenograft (X-2524) (Figure 3G). Because the tumor volume growth is mild in the single mutant xenografts X-3077 and X-3078 the volume difference between the initial tumor and after 10 days of treatment with BYL-719 is not as high as in dual mutant X-2524. Also, we noticed that BYL-719 treatment combined with LJM716, an anti-HER3 monoclonal antibody, is more effective in reducing tumor volume than BYL-719 treatment alone (Figure 3H). Dual mutation E726/H1047 makes the tumor grow significantly faster compared to the single mutant case. The double mutant tumor is also more sensitive to PI3K inhibitors.

However, not all doublets increase the PIK3CA oncogenic activity. For example, the impact of double mutation R88/T1025 (a combination of weak drivers) differs from E726K/H1047R in the screened PDX tumors. The growth rate of the tumor with R88/T1025 is slower than the tumor having a single mutation (at position 88). The tumor with only R88 is more responsive to PI3K inhibitors compared to that with R88/T1025 (Figure S5A-H).

### Linking Double Mutations to Clinical Data Using Cancer Cell Lines and Xenografts

Dual mutations may increase the activation strength and enhance drug response. In Figure 4 we show driver mutations combined with weak drivers or strong latent drivers in EGFR, BRAF, APC and PTEN. Below, we probe PTEN, APC and EGFR double mutations with respect to drug treatments.

**Figure 4.**
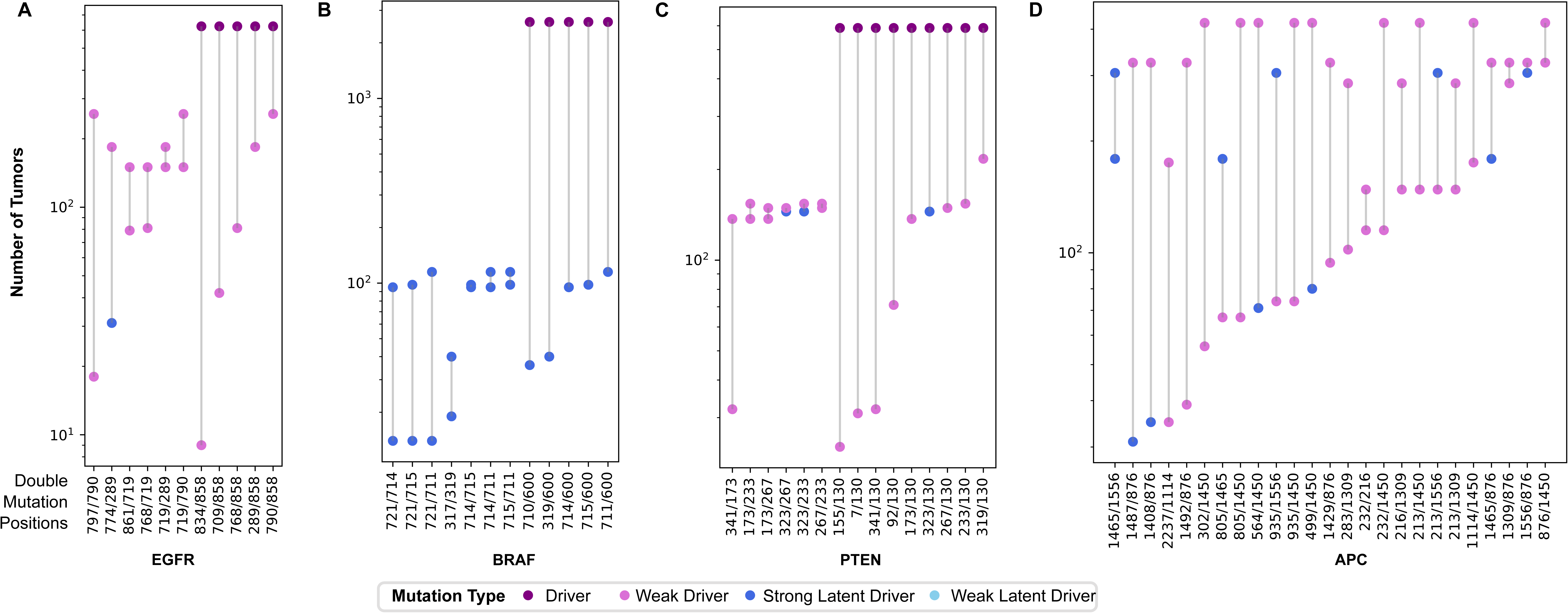
Representation of double mutations in EGFR, BRAF, APC and PTEN. Each paired dot represents one double mutation. Dots are colored according to their type, driver (purple), weak driver (orchid), strong latent driver (blue), weak latent driver (sky blue). **(A)** Double mutations and corresponding number of mutated tumors of each component reveals that there are 5 driver/driver, 5 weak driver/weak driver combinations and 1 weak driver/strong latent driver combinations; V774 might be a strong latent driver**. (B)** Genes harboring 12 double mutations, 7 are different combinations of strong latent drivers E317, E319, I710, L711, I714, E715 and L721, and the rest are V600 (driver) composed of a double mutation with the strong latent drivers. **(C)** PTEN carries 15 double mutations with only one potential strong latent driver (N323**). (D)** There are 25 double mutations on APC, R499, R564, E1408, S1465, T1487, T1556 that are potential strong latent drivers.

We screened all significant doublets across cell lines and PDX tumors. Double mutations are rare in the patient tumor samples and in cancer cell lines. Treatment data of patients are limited. Therefore, we aim to associate each marker double mutation with the cell lines or PDXs and assess their phenotypic impact through drug response data compared to their single mutant counterparts. In this way, we can assess the clinical impact of the same gene dual mutations and link the dual mutation patterns to drug response. We used the DepMap dataset together with Cell Model Passports to retrieve the mutational profile of cancer cell lines and the response to a panel of hundreds of drugs. We notice the same pattern: despite scanning hundreds of cancer cell lines, double mutations on the same gene are rare. In Figure 5, double mutations are linked to cell lines having the same pair, and cell lines are linked to drugs causing a significant response. These links are represented as a network of mutations, associated cell lines, and drugs. We listed some of the striking results on how dual mutations can alter the response to the drug in the same tissue.

**Figure 5.**
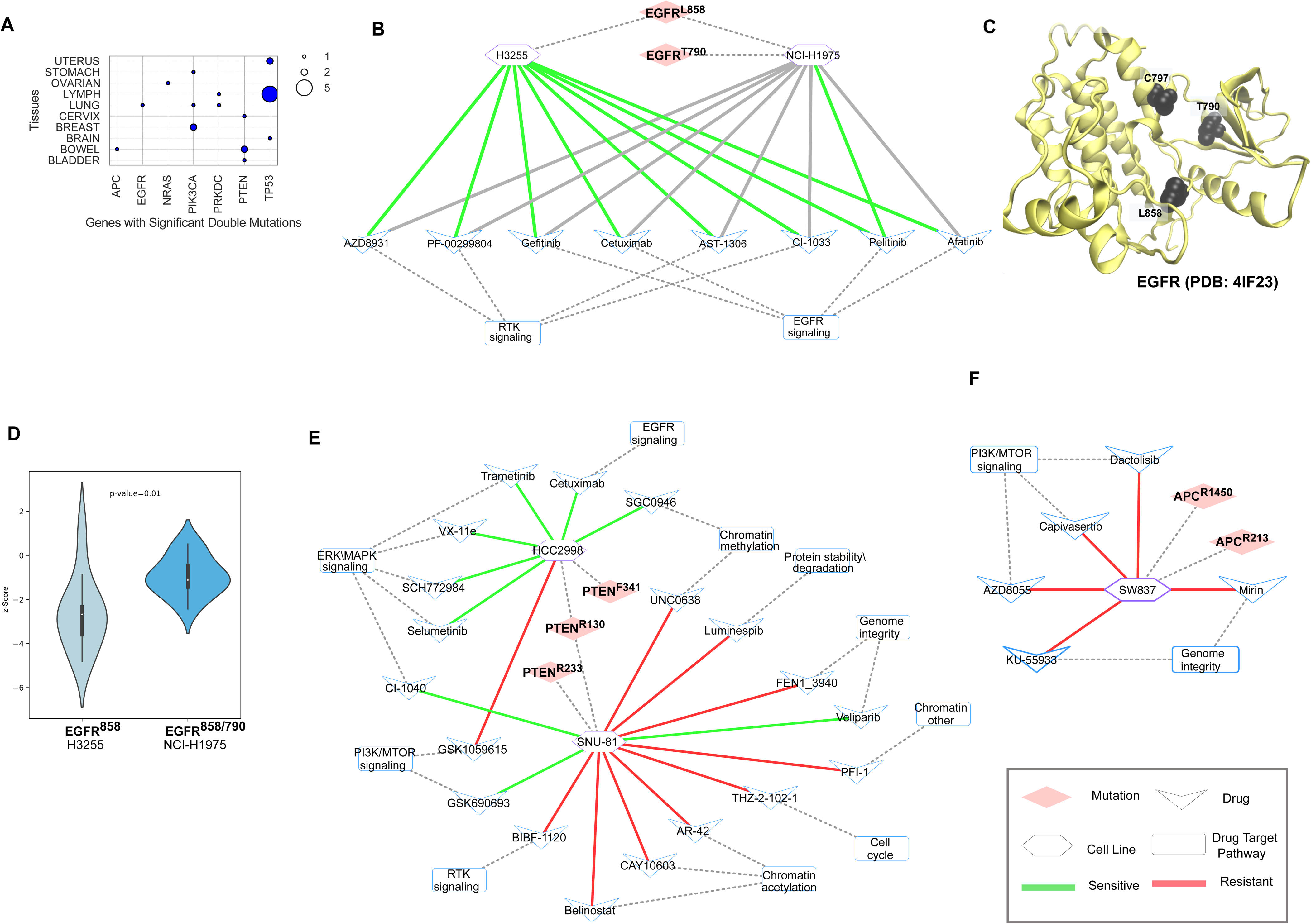
A wider analysis of double mutations in cell lines and association of doublets to drug response for clinical implications. **(A)** Prevalence of the double mutations in tissues associated with cell lines. APC, EGFR, NRAS double mutant cell lines belong to bowel, lung and ovarian tissues respectively. There are two cell lines in bowel tissue carrying PTEN double mutations, and five cell lines in lymph tissue with TP53 double mutations. **(B)** EGFR mutation doublets in lung cancer cell lines and their response to drugs in network representation. Among eight drugs targeting EGFR and RTK pathways, EGFR^L858/T790^ mutant cell line is only sensitive to Pelitinib, and it is unresponsive to the rest, although the EGFR^L858^ mutant cell line is sensitive to all. **(C)** Representation of dual mutations in EGFR structure. **(D)** EGFR mutation doublets in lung cancer together with the violin plot that shows the response to RTK inhibitor in dual mutant and single mutant cell lines. More negative z-score means more sensitivity and more positive z-score means more resistance to the drug molecule (**E)** PTEN and **(F)** APC mutation doublets in colon cancer cell lines and their response to drugs in network representation.

Among 228 same gene mutations only 17 doublets match with one or more cell lines in Cell Model Passports [36]. We constructed a network by using Cytoscape [50] with 4 different node types, mutations (components of a double mutation), cell lines, drugs, and drug target pathways. In the same gene double mutation network, there are 22 mutations, 19 cell lines, 206 drugs, and 22 drug target pathways nodes with 548 edges between them. In Figure 5A, we show the prevalence of the same gene dual mutations in corresponding tissues of the cell lines. These are consistent with the patient tumor doublets obtained in the previous section.

One example is EGFR^L858/T790^ doublet, a combination of two driver mutations (Figure 4A), in one cell line (NCI-H1975) of lung cancer. H3255 cell line has only one mutation at position L858 in EGFR (Figure 5B). Both mutations are on the tyrosine kinase domain to which the RTK inhibitors bind (PDB: 4IF23, Figure 5C). However, response to the inhibitors is significantly different in cell line with dual mutant EGFR which is more resistant compared to the single mutant cell line (p-value=0.01, Figure 5D).

BRAF has multiple double mutations (Fig. 4B). The most significant doubles contain the V600 strong driver and one of the strong latent drivers (I710, L711, I714, E715, L721). E715 is in the interface of the BRAF homodimer and the other strong latent drivers are in the same 3D cluster with E715. These mutations are not annotated based on their clinical or kinase activity; however, contributing to a significant double mutation make them strong latent driver candidates. They can be further analyzed based on their mechanism of action complementing V600. The mechanisms of BRAF mutations were classified into those signaling as active monomers, those acting as constitutive active dimers, and those having impaired/dead kinase activity [51]. Despite being very rare in our dataset, we have three cases of impaired/dead mutations paired with other classes: BRAF^G469/K601^, BRAF^G466/L597^, and BRAF^G466/V600^. Another rare double mutation is BRAF^L597/K601^. The double mutation components lead to a constitutively active dimer, independent of Ras. The mutations may increase Raf affinity. A combination of strong latent drivers located at or close to the dimer interface can fit into this activation mechanism. To identify these pairs a larger dataset is necessary, but still we can identify some rare doubles that need clinical evaluation.

A second example involves PTEN (Figure 5E). PTEN^R130/R233^ and PTEN^R130/F341^ doublets, both composed of a driver and a weak driver mutation (Figure 4C), exist in two cell lines which differ in response to drugs as compared to their single mutant counterparts. Although single mutant PTEN is resistant to drugs targeting ERK/MAPK signaling, cell lines having dual mutant PTEN are sensitive to these drugs. Additionally, cell line SNU-81 having PTEN^R130/R233^ becomes resistant to genome targeting drugs compared to single mutant cell lines.

The lower panel of Figure 5F presents APC dual mutations with associated cell lines. The subnetwork shows that cell line SW837 carrying dual mutant APC (R1450*/R213*, a combination of weak drivers (Figure 4D) becomes resistant to drugs targeting PI3K/TOR signaling when compared to the single mutant cell lines. Additionally, in the patient-derived xenograft dataset from [38], there is one xenograft carrying R1450*/R876* dual mutation (a combination of two weak driver mutations), model id X-1290, and one carrying single R1450* mutation, model id X-1173. We compared volume changes of these xenografts. For the dual mutant xenograft X-1290 is ∼1500 mm^3^ in the first 30 days and for the single mutant xenograft X-1173 ∼1200 mm^3^. The dual mutant xenograft does not encounter any tumor volume change during the first 70 days when treated with Cetuximab, an EGFR inhibitor, while the single mutant xenograft keeps growing around 900 mm^3^ during the first 15 days (Figure S6A-D).

Despite the small number of patients with follow-up information, Kaplan-Meier survival analysis comparing single and same gene double mutant patient groups demonstrates a significant difference between the two groups. Patients with double mutant PIK3CA (H1047/R88) and APC (R876/T1556) have worse survival than their single mutant counterparts. On the other hand, the patient group with PTEN double mutation (R130/R173) has a better survival than the single mutant group (Figure S7A-C).

## Discussion

The highly heterogeneous molecular profiles of tumors compel comprehensive studies to reveal the underlying patterns resulting in their dramatic phenotypic differences. Distinguishing cancer drivers from passengers has been one of the key objects of such studies. Here we scan the cancer genome landscapes aiming to identify latent drivers. We designed this study to discover latent driver mutations based on the hypothesis that some rare, or weak mutations can cooperate with other, *in cis* mutations, to enhance the oncogenic signal. In terms of population of conformations, conceptually, a strong driver mutation is close to a fully activated state with more than 90% of the population in the active state. However, the contribution of other types of drivers in the gene toward reaching a fully activated state can differ. For example, the contribution of a driver to the active state population can be between 50-75%, that of a weak driver is around 50% and the contribution of a strong latent driver is above 25%. This implies multiple mechanisms of action of double mutations relating to the combination of a driver mutation with either driver, weak driver, or strong latent driver. Latent driver mutations are protein context-specific having driver-like behavior but not identified as driver. We identified 228 significant, same gene double mutations which are composed of mostly one rare and one frequent mutation. Components of these double mutations are labelled as latent drivers if they have not been previously cataloged as driver. We newly cataloged 113 latent drivers. Despite being cancer-wide on their own, coupling with another mutation increases the cancer-type specificity and decreases the prevalence of these double mutations. The mutation load of tumors having a doublet in a tumor suppressor is significantly higher than in an oncogene, indicating their relative robustness to functional loss.

With the sparsity of patient treatment datasets, cell lines or patient-derived tumor xenografts are a useful clinical interpretation resource. We found significant differences in the response to PI3K inhibitors in double mutant PIK3CA samples which is in line with the recent work by Vasan et al [25, 27]. Additionally, tumor growth is extremely fast in double mutant PIK3CA compared to the single mutant. This phenotypic difference has been shown by Vasan et al for the couples of potential latent driver PIK3CA mutations E726, and weak drivers E453, M1043 with known driver mutations E542, E545, and H1047. Recent mechanistic studies suggest that the increased gene activity or acquired drug resistance is due to the mutation combinations. Zhang et al. [45] suggested that combinations of strong and weak drivers can enhance PI3K activity and explain the phenotypic differences in PIK3CA double mutant tumors, that we observed prominently in breast and uterus tumors. Here we further extended the analysis to combinations of less frequent mutations not catalogued as driver, which we view as potential latent drivers. Among them doubles with mutation at position R88 are depleted in breast but not in uterus, suggesting that potential latent driver mutations pairing with the mutation R88 are important signatures of uterus tumors.

Not limited to PIK3CA, numerous other significant double mutations with possible prognostic or therapeutic impact have also been identified (i.e. APC doublets in the bowel, EGFR in the lung in line with previous studies [26]). Some have not been previously analyzed clinically but have potential impact on drug response. For example, APC R1450/R876 double mutation results in significant sensitivity to cetuximab compared to single mutant APC R1450 in PDXs. On the other hand, cell lines having APC R1450/R213 doublet became resistant to PI3K/mTOR signaling inhibitors. Our approach also identified several rare same gene doublets, like the ones on cohesion complex subunits STAG2, RAD21, SMC3. This protein complex is important for sister chromatid cohesion, chromosome segregation, DNA repair, genome organization, and gene expression. The STAG2 subunit of the complex is highly mutated in bladder and myeloid cancers, and LOF mutations on STAG2 are correlated with DNA damage [52–54].

The sensitivity or responsivity of a drug action to a targeted cancer depends on how much the tumor relies on the particular oncogene and the cellular pathway with which it is associated. In PIK3CA, a combination of a driver mutation with either driver, weak driver, or strong latent driver, particularly under different mechanism of actions, have a good therapeutic response.

## Conclusions

In conclusion, we developed a comprehensive approach to discover latent driver mutations. We integrated molecular profiles of more than 80K patient tumors, drug treatment data of cancer cell lines and PDXs from multiple sources to reveal associations between molecular alterations to discover latent co-occurring driver mutations in the same allele, in non-redundant pathways and metastatic patterns with the help of multiple informatics techniques and interpret them through their clinical impact. Our results, supported by drug response data of cell lines and patient-derived xenografts, and transcriptomic profiles of single and double mutant tumors, provide a strong background for therapeutic potentials of double mutations. We believe that the results of this study may form a basis for further experimental evaluation of molecular alterations to be exploited for therapeutic purposes across different cancer types. Mechanistically, the actions of *same gene* double mutations are more straightforward to interpret as compared to double mutations in different protein in independent pathways. How double mutations in independent pathways work is highly challenging to understand.

## List of abbreviations

PDX: Patient-derived xenograft
TCGA: The Cancer Genome Atlas
GENIE: Genomics Evidence Neoplasia Information Exchange
PDB: Protein Databank

## Declarations

### Ethics approval and consent to participate

Not applicable

### Consent for publication

Not applicable

### Availability of data and materials

The results shown here are in whole or part based upon data generated by the TCGA Research Network: https://www.cancer.gov/tcga. The authors would like to acknowledge the American Association for Cancer Research and its financial and material support in the development of the AACR Project GENIE registry, as well as members of the consortium for their commitment to data sharing. Interpretations are the responsibility of the study authors. The cell line data underlying the results presented in the study are available from GDSC in https://www.cancerrxgene.org/downloads, Cell Model Passports in https://cellmodelpassports.sanger.ac.uk/downloads, and The Cancer Dependency Map project in https://depmap.org/portal/download/. The PDX data underlying the results presented in the study are available in Gao et al [38].

### Authors’ contributions

Conceptualization: CJT, RN, NT

Data curation: BRY

Formal analysis: BRY, CJT, NT

Methodology: BRY, CJT, RN, NT

Project administration: NT

Supervision: NT

Visualization: BRY, NT

Writing – original draft: 632 BRY, CJT, RN, NT

Writing – review and editing: BRY, CJT, RN, NT

## Supplementary Text

### Annotation of Double Mutations

We used domain and gene ontology (GO) information from InterPro, a consortium database collecting information from member databases, to annotate same gene double mutations (https://www.ebi.ac.uk/interpro/). Therefore a mutation position may match with more than one InterPro id, when this is the case we preferred Pfam annotation. If a mutation does not match with any Interpro id, we labelled it as “No_Domain_Info”. We mapped the Interpro ID’s to GO Annotations related to biological process, molecular function and cellular component categories. Usually an Interpro ID matches with more than one GO annotation. We constructed binary combinations of these domains for each component of a double mutation and counted the double mutations related to domain annotation combinations (similar procedure was conducted for GO annotations).

To find out spatial closeness of same gene double mutations we use 3DHotspots [1] which identifies statistically significant mutations clustering in 3D protein structures. There are 943 clusters of 504 different genes. If two mutated residues that are containing a dual mutation belong to the same cluster, we consider this same gene dual mutation components are in close proximity. We used Interactome Insider to identify if the components of either same gene or different gene double are located in the same interface [2]. Besides the experimental data in PDB and predicted data in Interactome3D, it also contains the predicted interfaces with their in-house method. We used EnrichR to find the pathway annotation of the genes having co-occurring mutations [3].

We matched the sequence position of each component of the same gene doublets to their InterPro domain, if available [4]. As a result, we mapped 113 out of 228 doublets to at least one Interpro domain. In case of more than one matching domain, we picked the Pfam original one, if available. Among them, only one component has domain information for 22 double mutations. In 96 doublets, both components have no domain information. We obtained a total of 19 InterPro domains. Mutations without any domain information are labeled ‘No_Domain_Info’ (this label covers cases whether the mutation position does not match with any domain or it belongs to loop, hinge regions). As shown in Figure S1A, a large portion of the mutations is in a region with domain annotation. Both components of 56 dual mutations are in the same domain. Doublets with domain information are either located in flexible or hinge or disordered regions or located in different domains.

A similar approach is applied to find GO molecular function information for partner mutations of doublets (Figure S1B). Both components of 57 dual mutations match with at least one GO molecular function annotation. Neither component of 151 dual mutations matches any GO Annotation. In the remaining 20 dual mutations, only one component matches with a GO annotation. Protein kinase activity and ATP binding are two molecular functions that are the most frequent annotations covering ∼15% of the same gene dual mutations having GO annotation.

### Alterations in Chemical Properties of amino acids

In order to classify alterations with respect to chemical classes of amino acids before and after mutations, we prepared a file containing unique rows as follows “patient barcode| gene | residue number | AA before mutation | AA after mutation”. We excluded the cases where the final amino acid is a stop codon. We calculated the fraction of chemical alterations on 2189 oncogene and 2262 tumor suppressor alterations among all oncogene and tumor suppressor alterations respectively (determined with respect to mutation positions). The 9 categories we evaluated in our analysis are Polar-Hydrophobic, Charged-Polar, Hydrophobic-Hydrophobic, Hydrophobic-Polar, Hydrophobic-Charged, Polar-Charged, Polar-Polar, Charged-Hydrophobic, Charged-Charged.

### PIK3CA Stability Analysis via Dynamut Tool

Using the inactive state (PDB id: 4OVV) we calculated the folding free energy (ΔΔG) upon mutation using DynaMut [5] to assess the impact of single and double mutations on PIK3CA stability. Unsurprisingly, considering their diverse mechanism of action no clear trend is observed (Figure S8). For example, H1047R is a strong driver that promotes interaction with the membrane. It destabilization impact is minor (ΔΔG ≈ -0.5 kcal/mol). The impact of weak drivers R88Q and R93W is somewhat stronger (ΔΔG ≈ -1.5 kcal/mol and ΔΔG ≈ -1 kcal/mol, respectively). The effect of allosteric mutation D539R is also minor (ΔΔG ≈ -0.6 kcal/mol). Another strong driver E542K (ΔΔG ≈ 0.7 kcal/mol), stabilizes the protein like the weak drivers D350G (ΔΔG ≈ 0.5 kcal/mol) and E453Q (ΔΔG ≈ 0.3 kcal/mol). The most prominent stability changes occur when the strong driver H1047R cooperates with the allosteric mutation P539R (ΔΔG ≈ -2.3 kcal/mol) and the minor mutation P104L (ΔΔG ≈ -2.5 kcal/mol). These two dual mutations H1047R/P539R and H1047R/P104L destabilize the protein as do T1025A/R88Q (ΔΔG ≈ 0.7 kcal/mol) while T1025A and R88Q have a destabilizing effect.

**Figure S1.**
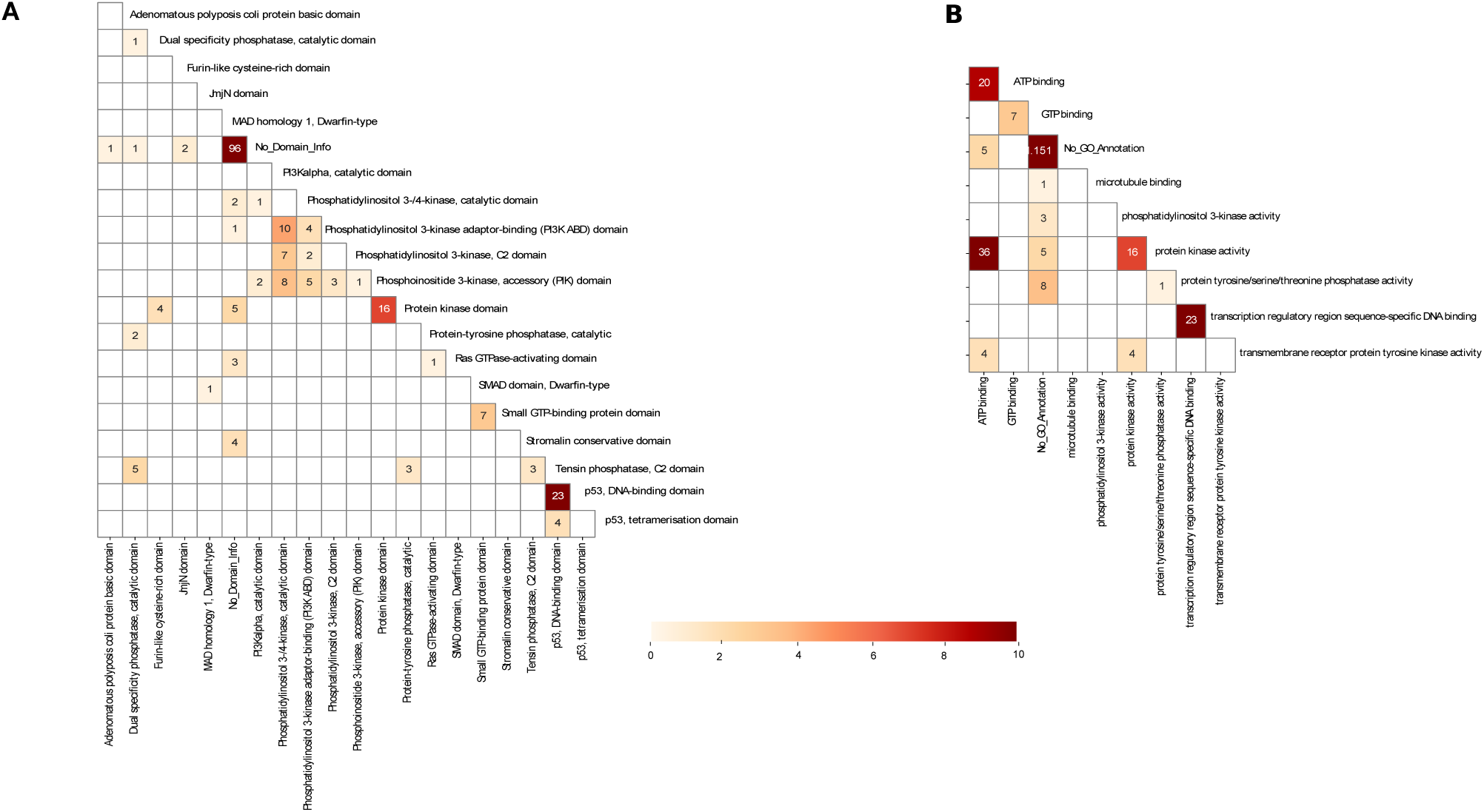
**(A)** Domain annotation and **(B)** GO molecular function annotation of the mutations in same gene dual mutations. The numbers in the squares correspond to the number of same gene dual mutations where constituents are from the domains on the x and y axes.

**Figure S2:**
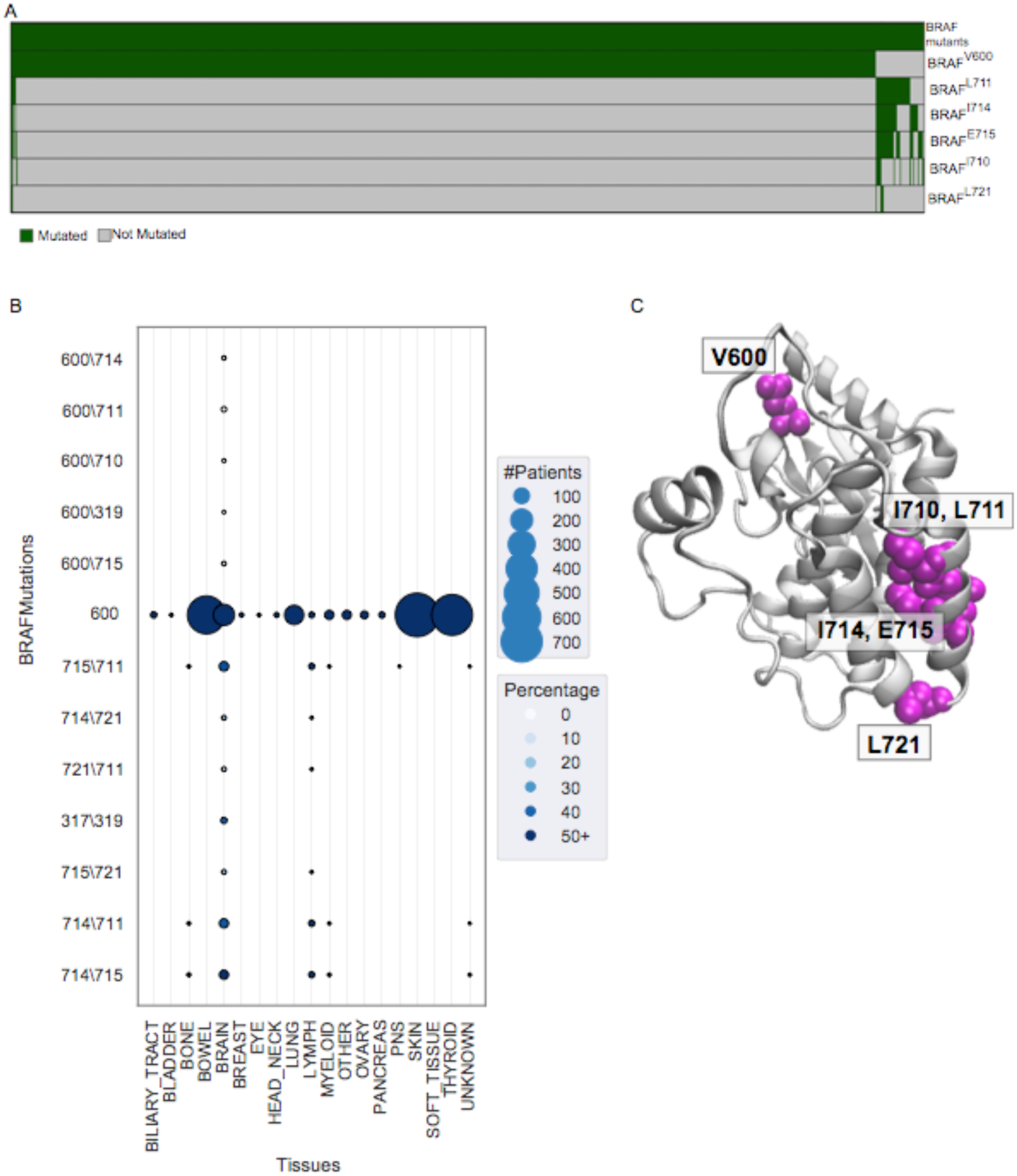
**(A)** Oncoprint of frequent BRAF dual mutation constituents. **(B)** Tissue prevalence of dual mutations of BRAF. **(C)** Mutations mapped to 3D structure of BRAF (PDB: 4G9R, Chain: B)

**Figure S3:**
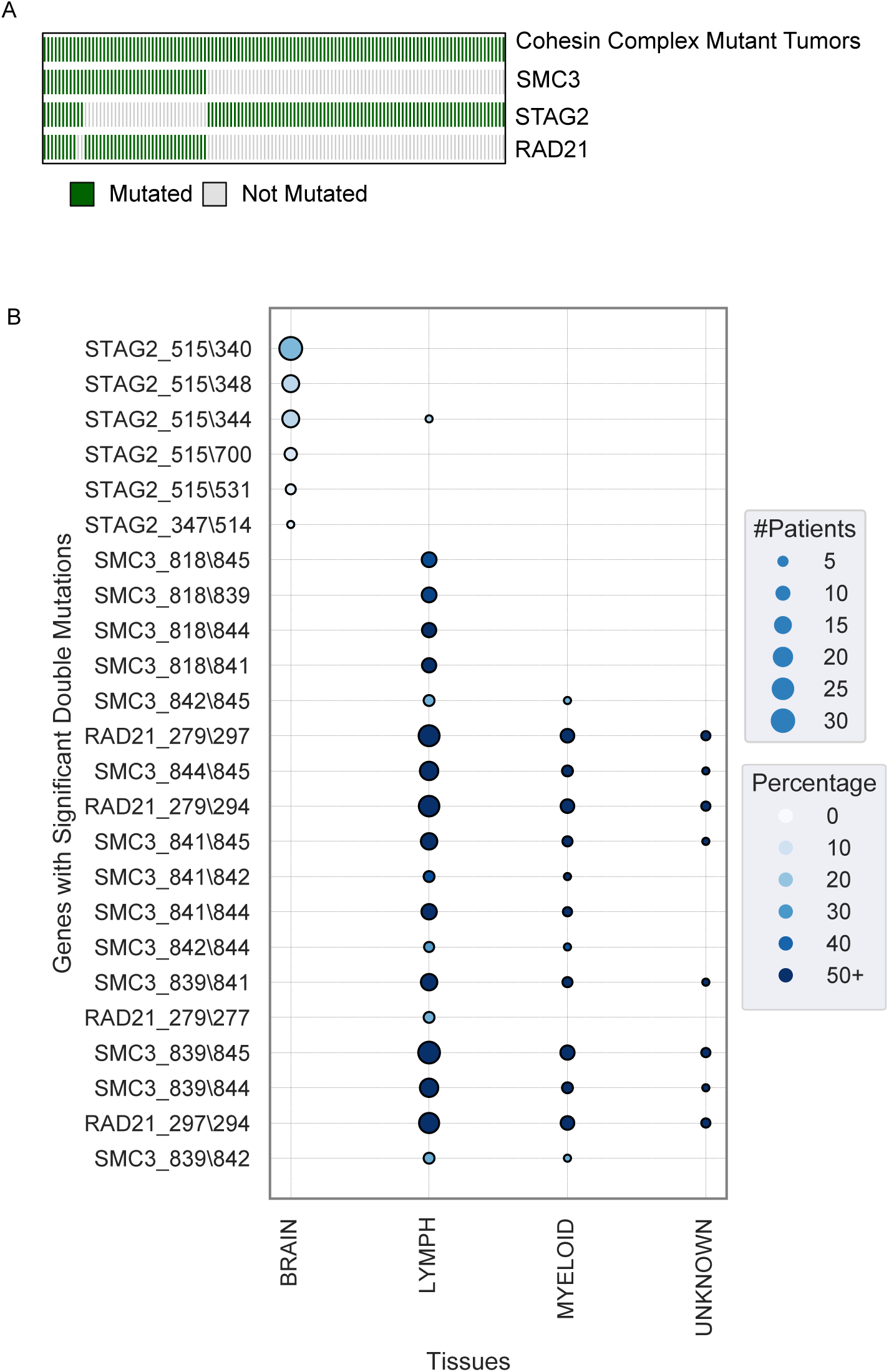
**(A)** Oncoprint of Cohesin complex subunits STAG2, RAD21, SMC3. **(B)** Tissue prevalence of dual mutations of STAG2, RAD21, SMC3.

**Figure S4:**
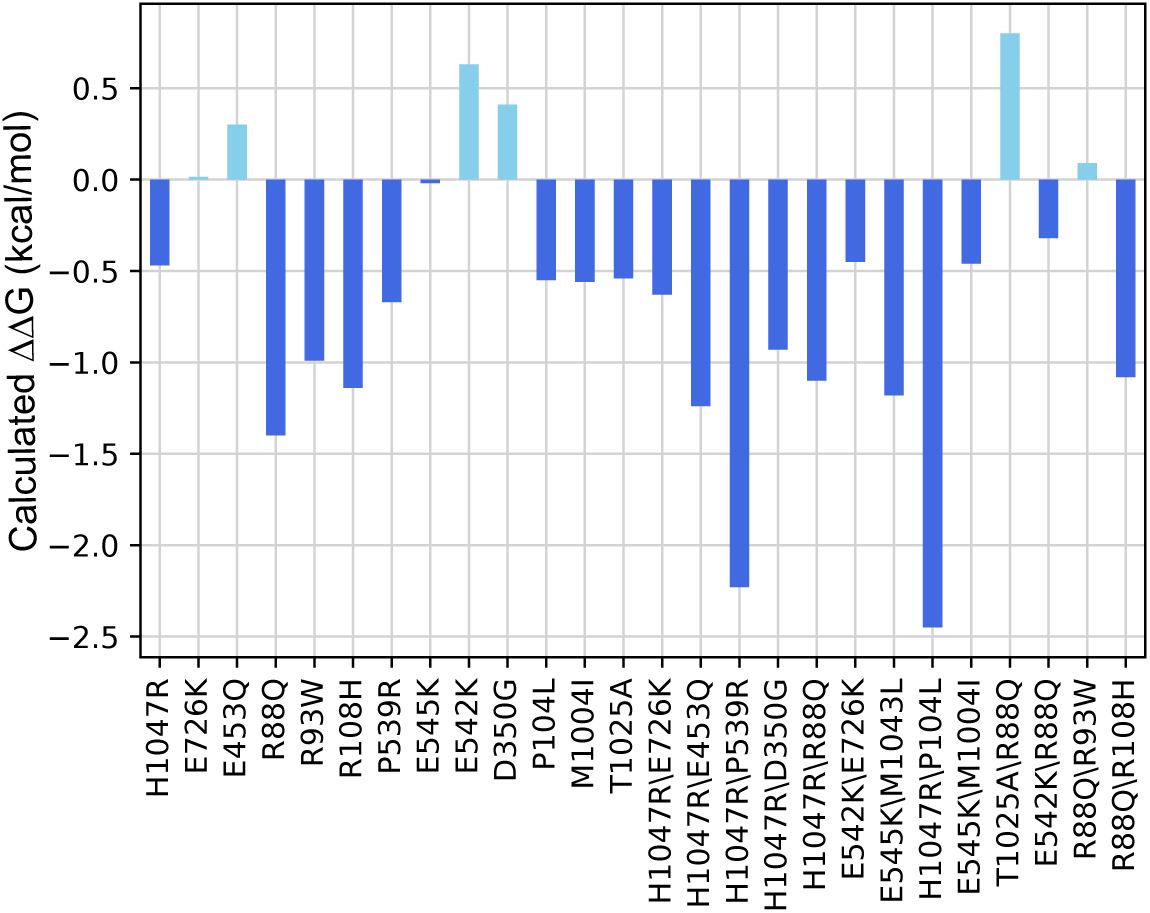
Predicted ΔΔG values for single and dual mutations of PIK3CA calculated with Dynamut web server (PDB id: 4OVV).

**Figure S5.**
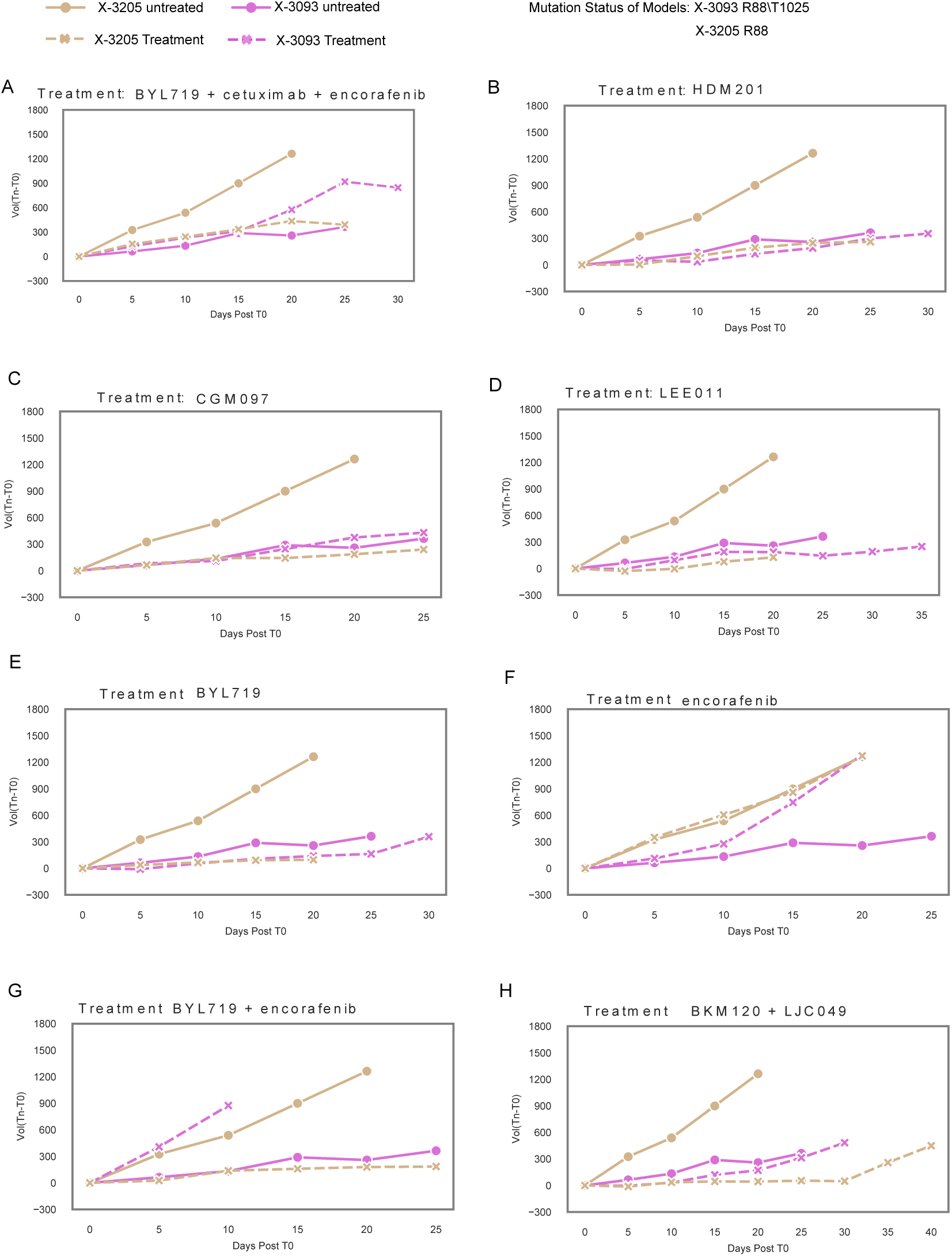
PIK3CA R88\T1025 mutant xenograft (X-3093, BRCA) volume change compared to single R88 mutant xenograft (X-3205, BRCA) for different drug treatments. x-axis shows treatment days, y-axis shows volume difference Volume(Day=n)-Volume(Day=0). Treatment with the drugs/drug combinations **(A)** BYL719+cetuximab+encorafenib combination. **(B)** HDM201 (Siremadlin). **(C)** CGM097. **(D)**LEE011(**Ribociclib**) **(E)** BYL719 (Alpelisib) **(F)** Encorafenib **(G)** BYL719+Encorafenib **(H)** BKM120+LJC049

**Figure S6.**
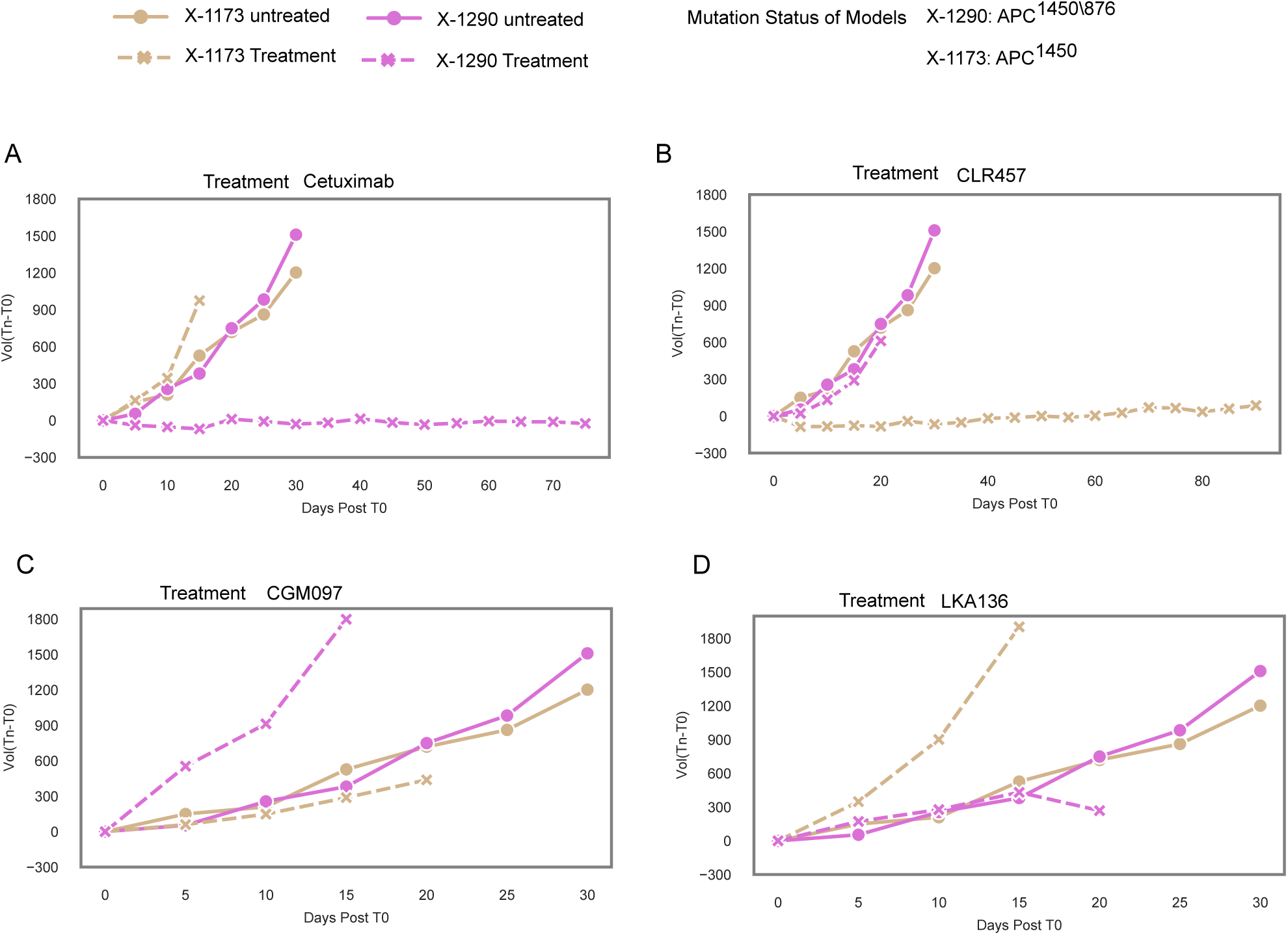
APC R1450\R876 mutant xenograft (X-1290, CRC) volume change compared to single R1450 mutant xenograft (X-1173, CRC) for different drug treatments. x-axis shows treatment days, y-axis shows volume difference Volume(Day=n)-Volume(Day=0). Treatment with the drugs **(A)** Cetuximab **(B)** CLR457 **(C)** CGM097 **(D)** LKA136.

**Figure S7.**
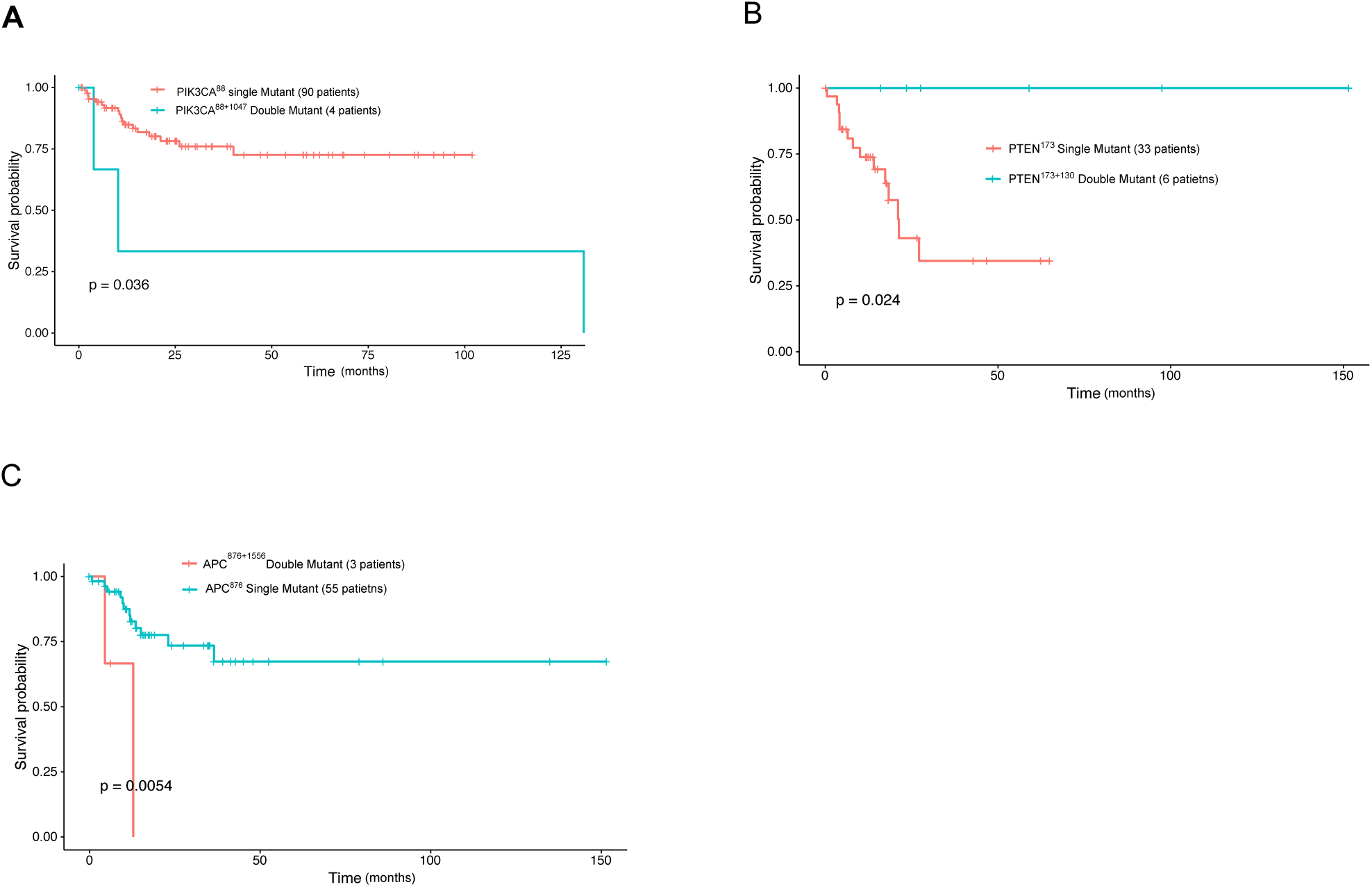
Kaplan-Meier survival analysis comparing single and double mutant patient groups. P-values are calculated with logrank test. **(A)** PIK3CA^88^ and PIK3CA^88+1047^ **(B)** PTEN^173^ and PTEN^173+130^ **(C)** APC^876^ and APC^876+1556^

